# Neural Processing of Taste-Related Signals in the Mediodorsal Thalamus of Mice

**DOI:** 10.1101/2024.08.05.606609

**Authors:** Katherine E. Odegaard, Cecilia G. Bouaichi, Greg Owanga, Roberto Vincis

## Abstract

Our consummatory decisions depend on the taste of food and the reward experienced while eating, which are processed through neural computations in interconnected brain areas. Although many gustatory regions of rodents have been explored, the mediodorsal nucleus of the thalamus (MD) remains understudied. The MD, a multimodal brain area connected with gustatory centers, is often studied for its role in processing associative and cognitive information and has been shown to represent intraorally-delivered chemosensory stimuli after strong retronasal odor-taste associations. Key questions remain about whether MD neurons can process taste quality independently of odor-taste associations and how they represent extraoral signals predicting rewarding and aversive gustatory outcomes. Here, using C57 male and female mice we present electrophysiological evidence demonstrating how MD neurons represent and encode 1) the identity and concentrations of basic taste qualities during active licking, and 2) auditory signals anticipating rewarding and aversive taste outcomes. Our data reveal that MD neurons can reliably and dynamically encode taste identity in a broadly tuned manner and taste concentrations with spiking activity positively and negatively correlated with stimulus intensity. Our data also show that MD can represent information related to predictive cues and their associated outcomes, regardless of whether the cue predicts a rewarding or aversive outcome. In summary, our findings suggest that the mediodorsal thalamus is integral to the taste pathway, as it can encode sensory-discriminative dimensions of tastants and participate in processing associative information essential for ingestive behaviors.

## Introduction

Our consummatory decisions depend on the taste of the food and the reward experienced while eating. Gustatory information is processed through neural computations that occur in interconnected brain areas (Vincis and Fontanini, 2019; Spector and Travers, 2005). The neural activity of taste-related hindbrain and forebrain regions has been shown to represent the chemosensory qualities and the hedonic value of gustatory stimuli and, in some cases, also accounts for sensory signals that anticipate food availability (Vincis and Fontanini, 2019). Although the contributions of many gustatory regions to taste-related processing have been explored, other areas, such as the mediodorsal nucleus of the thalamus (MD), may also play a role. Yet, our understanding of the MD’s involvement in taste processing remains limited. The MD is a multimodal thalamic nucleus involved in various cognitive and affective processes. MD neurons have been implicated in reward processing and play a role in associative learning, attention, and memory (Kawagoe et al., 2007; Oyoshi et al., 1996; Plailly et al., 2008; Parnaudeau et al., 2013). Based on these results and its connectivity to frontal cortical areas (Sherman, 2016), MD has been conceptualized as a higher-order thalamic relay. However, the MD may also process food-related chemosensory information crucial for ingestive behaviors. Anatomical evidence in rodents reveals that MD receives afferents from the parabrachial nucleus (PBN) (Krout and Loewy, 2000) - a key gustatory nucleus in the brainstem - and the primary olfactory cortical areas (Price and Slotnick, 1983), and MD is reciprocally connected to the GC (Gehrlach et al., 2020; Oh et al., 2014; Allen et al., 1991). Electrophysiological experiments have shown that MD neurons respond to orthonasally-delivered odors (Courtiol and Wilson, 2016). Furthermore, a recent study showed that, after rats were exposed to intraoral odor-taste mixtures for several days to establish strong retronasal odor-taste associations, MD neurons represented orally consumed odors, tastes and their mixtures (Fredericksen and Samuelsen, 2022).

Although these data implicate MD in the processing of food-related signals, key questions remain unanswered, especially with respect to taste-related information. First, while MD has been shown to be an integral component of the network processing intraoral odor-taste associative chemosensory signals, it remains to be determined whether MD neurons can process taste quality and concentration information without an established odor-taste association. Second, gustatory experiences are often perceived against the background of previous expectations. Signals from all sensory modalities frequently provide information predictive of general and/or specific taste outcomes and play a crucial role in shaping consummatory behaviors. How do MD neurons represent exteroceptive auditory cues that predict a rewarding and aversive gustatory outcome, and how does their response to the cue relate to their response to the predicted outcome?

To answer these questions, we collected recordings of the spiking activity of MD neurons in mice engaged in different behavioral tasks. First, we recorded the spiking activity of mice engaged in behavioral tasks in which they experienced 3 µl of different taste qualities or different concentrations of sweet and salty taste. To separate taste-evoked activity from electrophysiological correlates of licking, we trained mice to lick six times on a dry spout to receive each taste stimulus and did not start neural recording sessions until the taste-evoked licking rate was independent of stimulus quality and concentration. Second, we recorded MD neural activity after mice were conditioned to associate each of two distinct auditory signals with the availability of one of two tastants with opposite valence. In general, although the mediodorsal thalamus is usually not considered part of the taste pathway, our results imply that this perspective may need to be reevaluated. Our findings demonstrate that mediodorsal thalamus neurons actively encode both the identity and concentration of taste signals, and respond selectively to cues that predict rewarding and aversive taste outcomes relevant to making consummatory decisions.

## Methods

### Experimental design and statistical analysis

#### Data acquisition

The experiments in this study were performed on 22 wild type C57BL/6J adult mice (10-20 weeks old) that were purchased from The Jackson Laboratory (Bar Harbor, ME). Of these, 11 mice (6 females, 5 males) were used for electrophysiology experiments, and the remaining 11 were used for anatomical tracing experiments (retrograde tracing: 2 female, 4 males; anterograde tracing: 1 female, 4 males. Following arrival at the animal facility, mice were housed on a 12h/12h light-dark cycle with ad-libitum access to food and water. Experiments and training were performed during the light portion of the cycle. All experiments were reviewed and approved by the Florida State University Institutional Animal Care and Use Committee (IACUC) under protocol “PROTO202100006”.

### Surgical procedures

All animals were anesthetized with an intraperitoneal injection of a cocktail of ketamine (25 mg/ml) and dexmedetomidine (0.25 mg/ml). The depth of anesthesia was monitored regularly by visual inspection of breathing rate, whisker reflexes, and tail reflex. Body temperature was maintained at 35 °C using a heating pad (DC temperature control system, FHC, Bowdoin, ME). Once a surgical plane of anesthesia was achieved, the animal’s head was shaved, cleaned, and disinfected with iodine solution and 70% alcohol before positioning it in a stereotaxic plate.

For retrograde tracing experiments, a small craniotomy was drilled above the GC (AP: +1.2 mm, ML: ∼ 3.8 mm relative to bregma). A glass pipette was loaded with the cholera toxin subunit B conjugated to Alexa Fluor 555 (CTB-555; Thermo Fisher Scientific, Waltham, MA, catalog #: C34776) and lowered into GC (2.2 mm from the brain surface). We injected 150 nl of CTB-555 at a rate of 2 nl/s using a Nanoject III microinjection pump (Drummond Scientific, Broomall, PA). Following injection, we waited an additional five minutes before slowly extracting the glass pipette. Similarly, anterograde tracing was achieved by injecting the adeno-associated viral construct AAV9-hSyn-mCherry (2.4 × 10^13^ GC/ml; Addgene, Watertown, MA, catalog #: 114472-AAV9) in the GC at the following coordinates: AP: +1.5 mm, ML: ∼ 3.8 mm, DV: -2.5 mm and -2.2 mm, and AP: +1.3 mm, ML: ∼ 3.8 mm, DV: -2.6 mm and -2.2 mm). At each position, 300 nl of anterograde tracer was injected at 2 nl/s using the Nanoject III. We waited five minutes between each injection before moving to the next DV site or removing the pipette from the brain.

To record extracellular activity, mice were implanted with a chronic and movable silicon probe (P1, Cambridge Neurotech, Cambridge, UK) mounted on a nanodrive shuttle (Cambridge Neurotech). The probe had two shanks, each with 16 electrodes (organized in two adjacent rows spaced 22.5 µm apart), evenly spaced at 25-µm intervals. Craniotomies were opened above the left MD for implanting probes and above the visual cortex for implanting ground wires (A-M system, Sequim, WA, Cat. No. 781000). Unless specified otherwise, the probe was oriented with the shanks aligned rostro-caudally in the mouse brain. The anterior shanks of the P1 probes were positioned at AP: -1.3 mm and ML: +0.15 mm (relative to bregma) and slowly lowered above MD (2.7 mm below the cortical surface). The probes were further lowered 300 µm before the first day of recording of the experimental session. The probes and a head screw (for head restraint) were cemented to the skull with dental acrylic. Before implantation, the tips of the silicon probes were coated with a lipophilic fluorescent dye (DiI; Sigma-Aldrich, St. Louis, MO), which allowed visualization of probe locations at the end of each experiment. The animals were allowed to recover for a minimum of 7 days before beginning the water restriction regimen and training. The voltage signals from the probes were acquired, digitized, and bandpass filtered with the Plexon OmniPlex system (Plexon, Dallas, TX) (sampling rate: 40 kHz). The time stamps of task events (licking and stimulus delivery) were collected simultaneously through a MATLAB (MathWorks, Natick, MA) based behavioral acquisition system (BPOD, Sanworks, Rochester, NY) synchronized with the OmniPlex system.

#### Behavioral training and taste stimuli

One week prior to starting training, mice were mildly water restricted, receiving 1.5 ml of water per day and maintaining at least 80% of their pre-surgical weight. One week after starting the water restriction regimen, the mice became accustomed to being head restricted for short, 5-minute, daily sessions that gradually progressed over days to longer sessions. The body of the mouse was covered with a semicircular opaque plastic shelter during head restraint to limit body movements of the animal without stressful constriction. The fluid delivery system, licking detection, and behavioral paradigm have been described in detail in previous studies from our group (Bouaichi et al., 2023; Bouaichi and Vincis, 2020; Neese et al., 2022). Briefly, fluid stimuli were delivered by gravity using computer-controlled 12 V solenoid valves (Lee Company, Westbrook, CT) that were calibrated daily to deliver 3 µl from a licking spout made of short polyamide tubing (ID 0.03, MicroLumen, Oldsmar, FL). During experiments aimed at studying taste quality processing, mice were trained to lick the spout six times to trigger the delivery of a 3-µl drop of one of multiple taste stimuli including sucrose (0.1 M), NaCl (0.05 M), citric acid (0.01 M), quinine (0.001 M), and deionized water at room temperature. These stimuli represent a broad range of taste qualities and provide compatibility with prior studies of electrophysiology of tastes in awake mice (Bouaichi and Vincis, 2020; Bouaichi et al., 2023; Neese et al., 2022; Levitan et al., 2019; Dikecligil et al., 2020). Each trial involved the pseudo-random delivery of one of five gustatory stimuli (ensuring that each of the five stimuli appears once in every block of five trials), followed by a 3-µl water rinse administered 7 ± 1 s after the stimulus. Following rinse, an intertrial interval (ITI) of 6.5 ± 1.5 s separated consecutive trials. Deionized water was used both as a stimulus and a rinse. To study taste concentration, mice were similarly trained to lick the spout six times to trigger the delivery of a 3-µl drop of taste stimulus. For this experimental session, we focused on two taste qualities: sweet (sucrose) and salty (NaCl). The mice were trained during alternate daily sessions to experience different concentrations of NaCl (0.3 M, 0.15 M, 0.07 M, 0.04 M, and 0.02 M) and sucrose (0.3 M, 0.1 M, 0.06 M, 0.04 M, and 0.02 M). All tested concentrations are peri- and suprathreshold for mice (Treesukosol et al., 2009; Spector, 2015). Similar to what has been described above, each trial involved the pseudo-random delivery of one of five taste concentrations, followed by a 3-µl water rinse administered 7 ± 1 s after the stimulus. Following rinse, an ITI of 6.5 ± 1.5 s separated consecutive trials. For taste expectation experiments, mice were conditioned to associate each of two distinct auditory cues with the availability of one of two tastants (sucrose and quinine). For this task, we used higher concentrations of sucrose (0.3 M) and quinine (0.03 M) to facilitate learning. Auditory cues consisted of 2-second-long single tones (75 dB, 2 kHz; or 75 dB, 12 kHz). The start and end of auditory signals correlated with the start and end of the movement of the licking spout. Gustatory stimuli were delivered 500 ms after auditory cue offset. Cue-taste contingencies were counterbalanced between mice. Cue-taste pairings were presented in a block design, with ten pairings per block and at least six blocks per session. After at least five days of training, mice consistently licked in response to the sucrose-anticipating cue and strongly reduced anticipatory licks for the quinine-anticipating cue. Training ended when criterion performance was achieved (see below for more information on this analysis). All taste stimuli were purchased from Sigma-Aldrich and dissolved in deionized water to reach the final concentration.

#### Electrophysiology data acquisition and statistical analyses

Voltage signals from the probes were acquired, digitized and bandpass filtered with the Plexon OmniPlex system (Plexon, Dallas, TX) (sampling rate: 40 kHz). The time stamps of task events (licking and stimulus delivery) were collected simultaneously through a MATLAB (MathWorks, Natick, MA) based behavioral acquisition system (BPOD, Sanworks) synchronized with the OmniPlex system. Kilosort 3 (Pachitariu et al., 2016) was used for automated spike sorting on a workstation with an NVIDIA GPU, CUDA, and MATLAB installed. Following spike sorting, Phy software was used for manual curation. Finally, quality metrics and waveform properties were calculated using a code based on SpikeInterface (Buccino et al., 2020). Only units with an overall firing rate *>* 0.3 Hz, signal-to-noise ratio *>* 3.0, and an ISI violation rate *<* 0.2 % were used for the subsequent analyses. All of the following analyses were performed on custom MATLAB and R scripts. Across all experimental session we recorded a total of 536 neurons from 11 mice in 28 experimental sessions (16.7±5.2 neurons per session). Taste quality and concentration recording sessions were interleaved, and the probes were lowered _100 µm after each session. The spontaneous rate (calculated 5 s before stimulus) was 12.56 ±7.62 Hz while the evoked firing rate (calculated 5 s after stimulus) was 12.58 ±7.63 Hz. Based on established criteria for assessing bursting behavior (Lee et al., 2012; Guido et al., 1992), the majority of recorded neurons (86%) exhibited tonic firing patterns, suggesting a predominant mode of sustained activity in the population of MD neurons.

##### Taste selectivity

A neuron was defined as taste- or concentration-selective if 1) the evoked spiking activity differed significantly from the baseline activity (using “change point” analysis), and 2) it showed significantly different response profiles to the different stimuli (four tastants and water for taste quality experiments, the five different concentrations of sucrose and NaCl for the taste concentration experiments) using two-way ANOVA.

###### Change point analysis

Taste selectivity was first evaluated using a “change point” (CP) analysis (Jezzini et al., 2013; Liu and Fontanini, 2015; Vincis and Fontanini, 2016). Initially, we computed the cumulative distribution function (CDF) of spike occurrences in all trials for a given stimulus. For each neuron and stimulus, we analyzed the CDF within the time interval starting 1 s before and ending 2 s after taste delivery. Sudden changes in firing rates resulted in a piecewise change in the CDF slope, leading to the identification of a CP. If no CP was detected for any of the stimuli, the neuron was deemed non-responsive. If at least one significant CP was found after stimulus presentation, two-way ANOVA (see below) was performed to further establish taste selectivity.

###### two-way ANOVA

Stimulus selectivity was further assessed by evaluating differences in the magnitude or time-course of the taste-evoked firing rate across the five stimuli. We used a two-way ANOVA (with stimulus identity and time-course as factors) (Samuelsen et al., 2012; Jezzini et al., 2013; Liu and Fontanini, 2015; Bouaichi and Vincis, 2020) using 250-ms bins in the 0 to 1.5 s post-stimulus time interval. A neuron was considered taste-selective if the main effect of the stimulus identity or the interaction term (stimulus identity X time-course) was significant at p *<* 0.05. In addition, concentration-selective neurons were classified as exhibiting positive or negative monotonic responses based on their evoked firing rates at different concentrations of sucrose or NaCl. For each neuron, the average evoked firing rate was computed for the lowest and highest concentrations tested. If the firing rate at the highest concentration was greater than that at the lowest concentration, the neuron was classified as positive monotonic; conversely, if the firing rate at the highest concentration was lower than that at the lowest concentration, the neuron was classified as negative monotonic. This classification was applied separately for sucrose- and NaCl-responsive neurons.

###### Sharpness index and entropy

To further investigate the response profile of MD neurons, we used the sharpness index (SI) (Rainer et al., 1998; Yoshida and Katz, 2011) and the entropy (H) (Smith and Travers, 1979), two standard methods used to evaluate taste tuning. SI was calculated on the mean firing rate during the 1.5-s interval after taste delivery and was defined as:

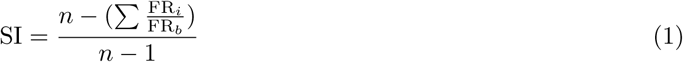

where FR*_i_* is the mean firing rate for each taste (*i* = 1 − 5), FR*_b_* is the maximum firing rate among gustatory stimuli, and *n* is the total number of stimuli (*n* = 5). An SI of 1 indicated that a neuron responded to one stimulus (narrow tuning), and the value 0 indicated equal responses across stimuli (broad tuning). Entropy metric H was computed as previously described by Smith and Travers (1979):

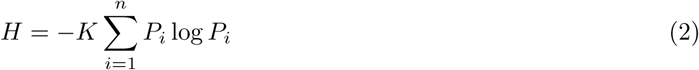

where *K* is a constant (1.43 for 5 stimuli) and *P* is the proportional response to each gustatory stimulus (*i*). Proportional taste responses were obtained by subtracting the mean taste-evoked firing rate (over 1.5 s after taste delivery) from the mean firing rate preceding taste (over 0.5 s before taste delivery). To control for negative proportional taste responses (that is, when taste-evoked firing rate *<* mean firing rate preceding taste; when the taste results in suppression of spiking activity), the absolute value of the proportional taste responses was included in the analysis. This is required because *logP_i_* is only defined for positive values of *P_i_*. Overall, a low *H* indicated a narrowly tuned taste-selective neuron, while a high *H* indicated a broadly tuned taste-selective neuron.

##### Selectivity index to track associative learning

Lick time stamps were aligned with the cue onset and PSTHs were constructed (bin size is 100 ms). ROC analysis (Vincis and Fontanini, 2016; Fonseca et al., 2018; Feierstein et al., 2006) was then used to compare the mean licking rates between the cue-Q and cue-S trials in one temporal epoch (1.5-2.5 s) during the training sessions, before the neural recording started. Specifically, the area under the ROC curve (auROC) was used to calculate the selectivity index as: selectivity index = 2 × (auROC-0.5). The selectivity index ranged from -1 to 1, where -1 means a higher licking rate in cue-S or sucrose trials, 1 means a higher licking rate in cue-Q or quinine trials, and 0 means a similar firing rate between the two cues or taste outcome trials. To assess the significance of the selectivity index, we used a permutation test where the trials were shuffled without replacement. Data were shuffled 1000 times and the pseudo-selectivity index was calculated for each iteration of the shuffling. The p-value was computed by comparing the actual selectivity index with the pseudo-index. We used a p *<* 0.01 criterion to determine significance.

##### Cue and taste/outcome selectivity

For the experiment investigating how MD neurons process taste-specific expectation, the stimulus selectivity was calculated as follows. Similarly to what we described above, a neuron was defined as selective for cues or taste outcomes if 1) evoked spiking activity differed significantly from baseline activity (using “change point” analysis), and 2) it showed significantly different response profiles to the two auditory cues (for cue selectivity) or to the two taste outcomes (for taste outcome selectivity) using two-way ANOVA.

###### Change point analysis

Cue and taste outcome-selectivity was first evaluated using CP analysis. As described above, we computed the CDF of spike occurrences in all trials for a given auditory cue or taste outcome. For each neuron and stimulus, we analyzed the CDF within a given time interval (for cue selectivity, starting 0.5 s before and ending 2.5 s after cue onset; for taste outcome selectivity, starting 0.5 s before and ending 2.5 s after taste outcome delivery). Sudden changes in firing rates resulted in a piecewise change in the CDF slope, leading to the identification of a CP. If no CP was detected for any of the stimuli, the neuron was deemed non-responsive. If at least one significant CP was found after stimulus presentation, two-way ANOVA (see below) was performed to further establish cue selectivity and/or taste outcome selectivity.

###### two-way ANOVA

Stimulus selectivity was further assessed by evaluating differences in the magnitude or time-course of the stimulus-evoked firing rate in the two cues or the two taste outcomes. We used a two-way ANOVA (with stimulus identity and time-course as factors) (Samuelsen et al., 2012; Jezzini et al., 2013; Liu and Fontanini, 2015; Bouaichi and Vincis, 2020) using 500-ms bins in the 0 to 2.5 s post-stimulus time interval. A neuron was considered selective if the main effect of the stimulus identity or interaction term (stimulus identity X time-course) was significant at p *<* 0.05.

##### Population decoding

To understand how well MD encodes information about the identity of gustatory stimuli and how taste information is processed over time, we used a population decoding approach (Meyers, 2013). To do this, we first constructed a pseudo-population of MD neurons using taste-selective neurons recorded in different sessions (taste quality: n = 141; taste concentration, sucrose: n= 87; taste concentration,

NaCl: n= 39) and using cue-selective neurons for taste-specific expectation sessions (n = 47). We then generated a firing rate matrix (trials X time-bin) where the spike time stamps of each neuron (1 s before and 1.5 s after taste) were re-aligned to taste delivery or auditory cue onset (for taste expectation), binned into 150-ms time bins with a 50-ms moving window, and normalized to Z-score. To assess the amount of stimulus-related information, we used a “max correlation coefficient” classifier. The spike activity data contained in our matrix were divided into cross-validation “splits” such that 80% was used for training (i.e., used by the classifier algorithm to “learn” the relationship between the population’s neural activity pattern and the different stimuli) and 20% for testing (i.e., predict which stimulus was delivered given the population’s pattern of activity that was used to train the classifier). This split was done to ensure that the model was trained on a substantial portion of the data while retaining a separate and independent subset for evaluation. This process was repeated 50 times (i.e., resample runs) per cross-validation split to calculate the decoding accuracy, defined as the fraction of trials in which the classifier made correct stimulus predictions. Comparisons of the classification accuracy between real and shuffled data or between different populations were performed using a permutation test (Ojala and Garriga, 2010).

##### Histological staining

At the end of the experiment, mice were terminally anesthetized and transcardially perfused with 30 ml of PBS followed by 30 ml of 4% paraformaldehyde (PFA). The brains were extracted and postfixed with PFA for 24 h, after which coronal brain slices (100-µm thick) were sectioned with a vibratome (VT1000 S; Leica, Wetzlar, Germany). To visualize the anatomical tracers and the tracks of the probes, brain slices were counterstained with Hoechst 33342 (1:5,000 dilution, H3570; ThermoFisher, Waltham, MA) using standard techniques and mounted on glass slides. Brain sections were viewed and imaged on a fluorescence microscope or with an Olympus FV1000 laser scanning confocal microscope. For the quantification of retrogradely labeled neurons, only Z-stack images acquired with the Olympus FV1000 confocal microscope were used. The use of Z-stacks allowed for better discrimination of individual neurons from surrounding neuropil, ensuring more accurate counts. Retrograde CTB^+^ neurons were manually counted with the Fiji Cell Counter plugin on four Z stacks in the same image plane for each thalamic nucleus (MD and VPMpc). The proportion of CTB^+^ neurons within each thalamic nucleus was calculated with the formula [(number of CTB^+^ neurons within a thalamic nucleus) / (total number of CTB^+^ neurons across all thalamic nuclei)]. This normalization accounts for variations between mice in the efficiency of injection and CTB labeling.

## Results

### MD reciprocal connection with the gustatory insular cortex

To investigate the type of taste-related information processed by mediodorsal thalamus (MD) neurons in mice, we first evaluated the neural connection between the primary taste cortex (i.e. the gustatory insula, GC) and MD using anatomical tracing approaches. The rationale for these experiments was twofold: 1) to confirm and expand on the existing evidence of the connection between the gustatory portion of the insular cortex (gustatory cortex, GC) and the MD in rodents (Allen et al., 1991; Gehrlach et al., 2020); 2) to locate the specific parts of the mouse MD that could potentially be involved in processing taste and/or taste-related information. Given the small size of this thalamic nucleus in mice, the latter step was crucial for us to ensure accurate targeting for silicon probe implantation for neural recording, the focus of the remainder of this study.

For these experiments, we first injected a retrograde neural tracer (cholera toxin subunit B conjugated with Alexa Fluor, CTB-555; Fig. 1A-B) into a region of the GC known to process gustatory information (Chen et al., 2011; Bouaichi and Vincis, 2020; Levitan et al., 2019; Bouaichi et al., 2023; Kusumoto-Yoshida et al., 2015) and receive direct axonal projections from the gustatory thalamus (ventroposteromedial parvicellular nucleus; VPMpc)(Chen et al., 2011; Fletcher et al., 2017; Chen et al., 2021). This approach allowed us to label the somata of neurons with terminal fields in the GC (Fig. 1A-B). The injections were unilateral and targeted all subdivisions of the GC (granular, dysgranular, and agranular; Fig. 1A, *left panel*). As anticipated, we observed a substantial number of thalamocortical neurons labeled in the ipsilateral VPMpc (Fig. 1A, *middle panel*). Further, beyond the VPMpc, we identified neurons labeled in the MD (Fig. 1A, *right panel*). To evaluate the relative distribution of GC-projecting neurons, we compared the number of CTB^+^ neurons in the two thalamic nuclei. Analysis of the relative number of retrogradely labeled neurons revealed significant differences in the distribution of thalamo-GC projecting neurons across the two nuclei (Figure 1B; Wilcoxon rank sum, W = 16, p-value = 0.02857). While the majority of labeled neurons originated from the gustatory thalamus (VPMpc), the presence of a substantial population in MD suggests that this higher-order thalamic nucleus may also play a role in influencing GC activity. Figure 1C shows the qualitative evaluation of the location of the MD neurons that projected to the GC obtained from six mice, with the largest overlap present in the medial subregion of the MD. To further explore this neural connection, we then used anterograde viral tracers (AAV9-hSyn-mCherry) in the GC to qualitatively map the projections from the GC to the MD (Fig. 1C). Similar to what was observed for retrograde tracer experiments, the medial subregion of the MD appeared to be the recipient of most of the GC-MD projections (Fig. 1D). Although MD is not part of the classical taste sensory pathway, its reciprocal connection with the GC suggests a potential role in processing gustatory information even in the absence of a previously established association with another oral sensory modality, such as retronasal odor (Fredericksen and Samuelsen, 2022). For these reasons, we next investigated the response profile of MD neurons in representing chemosensory gustatory information, focusing on taste quality and taste concentration, in active licking mice.

**Figure 1.**
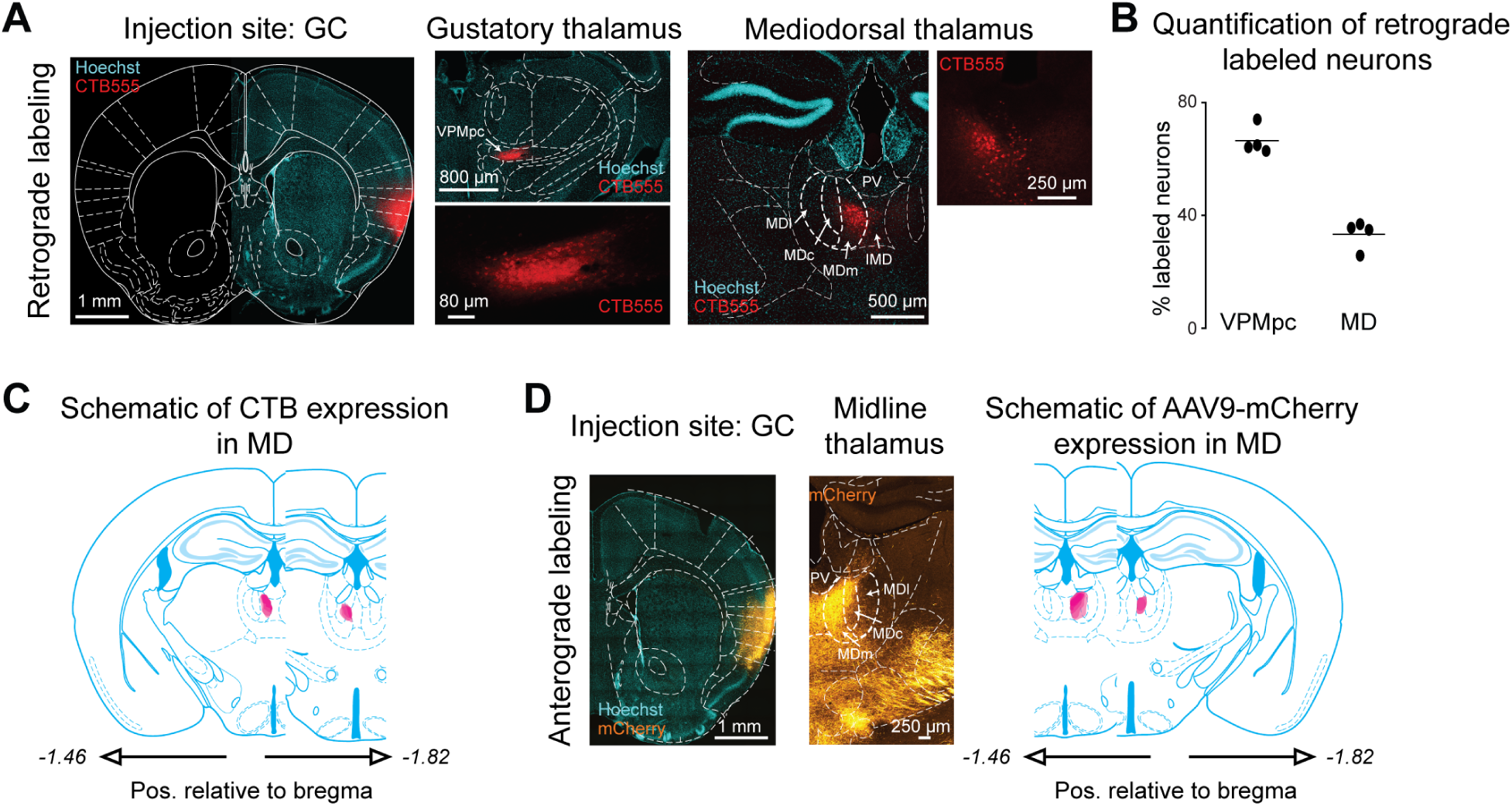
(A) Leftmost panel: coronal section of the mouse brain showing the CTB injection site (in red) in the GC counterstained with Hoechst (in cyan). Central panel: on the top, a coronal section showing the location of CTB^+^ neurons in the gustatory thalamus (VPMpc); on the bottom, high magnification image of CTB^+^ neurons in the VPMpc. Rightmost panel: a coronal section showing the location of CTB^+^ neurons in the MD with nuclear Hoechst counterstaining in cyan. On the right, high magnification image of CTB^+^ neurons in the MD. (B) Individual percentages (filled circles) and averaged (horizontal bars) (n = 4 mice) of CTB^+^ neurons in the two thalamic nuclei. (C) Qualitative reconstruction of the MD regions showing the highest amount of CTB^+^ neurons (n = 6 mice). (D) Leftmost panel: coronal section of the mouse brain showing the injection site of the adeno-associated viral construct AAV9-hSyn-mCherry (in orange) in the GC counterstained with Hoechst (in cyan). Central panel: a coronal section showing the location of mCherry^+^ axonal projections in the midline thalamic nuclei. The white dotted lines highlight the MD borders. Rightmost panel: a qualitative reconstruction of the MD regions showing the highest amount of mCherry^+^ axonal projections (n = 5 mice). Note that the qualitative labeling schematics shown in C and D refer only to MD and not to adjacent areas. MD: mediodorsal nucleus of the thalamus; MDc: mediodorsal nucleus of thalamus, central division; MDl: mediodorsal nucleus of thalamus, lateral division; MDm mediodorsal nucleus of thalamus, medial division; PV: paraventricular nucleus of thalamus; IMD: intermediodorsal nucleus of thalamus; VPMpc: parvicellular portion of the ventroposteromedial nucleus, also known as gustatory thalamus.

### Taste quality processing in the mouse MD

To investigate how individual MD neurons represent taste quality information in freely licking mice, we recorded single units using movable silicon probes (Cambridge Neurotech) mounted on a nanodrive shuttle (Cambridge Neurotech) implanted unilaterally in the MD (Fig. 2A-B). After habituation to head restraint, water-deprived mice were engaged in a behavioral task in which they had to lick six times on a dry spout to obtain a 3-µl drop of one out of four gustatory stimuli at a fixed concentration (0.1 M sucrose, 0.05 M NaCl, 0.01 M citric acid, 0.001 M quinine) or deionized water, all presented at room temperature (Fig. 2C). To distinguish neural activity evoked by gustatory stimuli from electrophysiological correlates of sensory and oromotor/palatability aspects of licking, we initiated neural recording sessions only after confirming that the licking patterns evoked by each gustatory stimulus were statistically similar within a 1.5-s post-stimulus temporal window, as assessed using a Kruskal-Wallis test (Bouaichi et al., 2023). The taste quality-independent licking ensured that differences in neural responses between gustatory stimuli within 1.5 s after taste delivery could not be attributed to overt licking/palatability behavioral correlates. Figure 2D shows the raster plots and PSTHs of three representative MD neurons. Visual inspection of the graphs indicated that each of these neurons responded to more than one taste stimulus with time-varying and multi-phasic changes in activity similar to those reported by Fredericksen and Samuelsen (2022) when oral stimuli were delivered via intra-oral cannulae. As a first step, we wanted to understand how many MD neurons, such as the three shown in Figure 2D, were selectively modulated by gustatory stimuli. Neurons were defined as taste-selective if they exhibited a significant change in activity compared to baseline for at least one taste stimulus and had significantly different responses to the five stimuli (see Methods). This analysis revealed that more than half of the MD neurons recorded (66.5%, 141/212) were taste-selective (Fig. 2E; *χ*^2^ test for given probabilities: *χ*^2^ = 23.113, df = 1, p-value = 1.527e-06).

**Figure 2.**
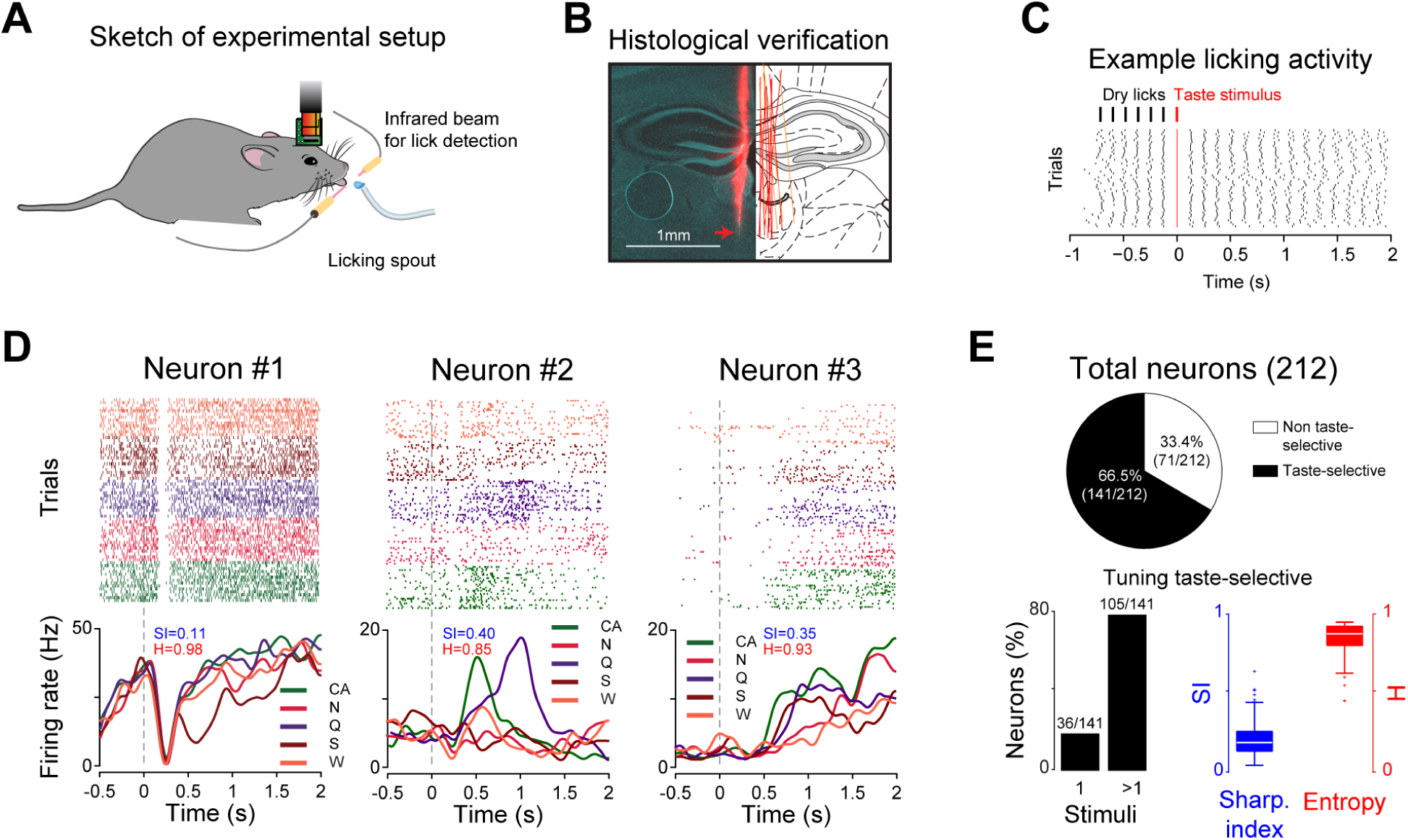
(A) Schematic showing the recording setup and a head-restrained mouse licking a spout to obtain tastants. (B) Left: example of histological sections showing the tracks (red) of 1 shank of the CN-P1 probe in the MD. Red arrow points to the tip of the probe. Right: schematic of the summary of probe tracks from the mice used in this study. Red shanks are implanted along the caudal-rostral axis (one shank visible), while yellow shanks are perpendicular, showing both shanks. (C) Top: diagram of the taste delivery paradigm: taste stimuli are delivered after six consecutive dry licks to the spout; bottom: raster plot of licking activity during one experimental session: each line represents an individual lick. Taste delivery occurs at time 0 s (highlighted in red). (D) Raster plots and PSTHs of three MD neurons showing taste responses. Trials pertaining to different tastants are grouped together (in the raster plots) and color-coded (both in the raster plots and PSTHs), with sucrose (S; *brown*), quinine (Q; *purple*), NaCl (N; *red*), citric acid (CA; *green*), and water (W; *orange*). Smaller text within the PSTHs highlights the sharpness (SI; blue) and entropy (H; red) values for each of the neurons. (E) Top: pie chart showing the proportion of MD taste-selective neurons. Bottom left: fraction of taste-selective neurons responding to one or more gustatory stimuli. Bottom right: the distribution of the breadth of tuning [expressed as sharpness index (SI; *blue*) and entropy (H; *red*)] of the taste-selective neurons. High SI or low H values indicate narrowly responsive neurons, whereas low SI or high H values imply that the same neuron is modulated by multiple tastants.

Next, we evaluated the tuning profile of the taste-selective neurons. We aimed to understand whether the taste-selective MD neurons preferentially responded to only one single taste stimulus (i.e., narrow tuning) or, as suggested by the example neurons shown in Figure 2D, if they were capable of encoding information pertaining to multiple tastes (i.e., broad tuning). This analysis served only to estimate how many taste-selective neurons were modulated by more than one tastant independent of the quality / identity of the taste. Our analysis revealed that, while only 25.5% (36/141) of MD taste-selective neurons responded to one taste, the vast majority (74.5%; 105/141) were modulated by more than one gustatory stimulus (Fig. 2E; *χ*^2^ test for given probabilities: *χ*^2^ = 33.766, df = 1, p-value = 6.216e-09). To further investigate differences in the tuning profiles of MD neurons, we calculated the response sharpness index (SI) and the response entropy (H) for each taste-selective neuron. These two analyses are standard techniques used to evaluate the breadth of tuning of single neurons (Smith and Travers, 1979; Rainer et al., 1998; Yoshida and Katz, 2011; Bouaichi and Vincis, 2020). Figure 2E shows the distribution of the response SI (Yoshida and Katz, 2011). An SI of 1 describes a neuron that responds to only one taste, and an SI of 0 describes a neuron that responds to all five stimuli. The distribution of SI values strongly implied that most taste-selective neurons were broadly tuned, suggesting that the majority of taste-selective neurons in MD are modulated by more than one taste. Similar results were obtained by analyzing the response H. Low H values are evidence of narrowly tuned neurons, whereas high H values indicate broadly tuned neurons. The results of this analysis further confirmed that MD neurons are broadly tuned.

After characterizing the profiles of chemosensory stimuli in individual MD neurons, we shifted our focus to the neural activity at the population level. While the activity of single neurons can represent important features of sensory stimuli, it is the information encoded in populations or ensembles (networks) of neurons that informs behavioral choices. As a result, we computed the taste quality decoding performances in our taste-selective neuron population (n = 141; gold trace in Fig. 3A). The onset of taste decoding occurred in the third temporal bin, approximately 200 ms after stimulus delivery, and reached its peak later than 500 ms after taste delivery (Fig. 3A). Additionally, while the overall classification value started decreasing after 500 ms, decoding performances remained significantly above chance and shuffled control (gray trace in Fig. 3A; permutation test, p *<* 0.05) until the end of the temporal window analyzed (Fig. 3A). As a control, the same analysis was performed using licking instead of spiking data (Fig. 3B). As expected, the taste classification accuracy using licking data never exceeded shuffled control level. The lack of taste classification with licking data, as opposed to spiking data, further indicated that the MD neural activity encodes specific information about the taste quality that could not be attributed to overt oromotor/palatability behavioral correlates. Figure 3C shows how classification during a 1.5-s post-stimulus temporal window changed as the decoder gained access to progressively more MD neurons. While the classifiers that were trained with a small number of neurons were less accurate at identifying taste quality information, increasing the number of neurons in the population drastically increased the decoding performance, reaching up to 50% accuracy with less than 20 neurons. Next, we wanted to provide detailed information on the temporal dynamics of the taste classification performance with respect to each gustatory stimulus. To do this, we constructed confusion matrices and characterized the classification performance for each tastant in 150-ms time-bins with 50-ms moving window around taste delivery (Fig. 3D). Figure 3A shows that the average decoding performance in taste-selective neurons was well above chance and shuffled control during the first 500 ms (between around 200 and 500 ms). However, inspection of the confusion matrices in Figure 3D revealed that, in this first phase of taste processing, not all taste qualities were equally classified. In this plot, the main diagonal represents the fraction of trials where the classifier accurately matched the predicted taste stimulus to its actual category. Comparison of the fraction of trials correctly classified in the first 500 ms revealed that all stimuli, with the exception of quinine and deionized water, were predicted by the decoding algorithm (Marascuilo’s test, Tables 1 and 2). As time progressed, all stimuli were decoded with accuracy above chance (20%) within 1 s from fluid delivery. However, as represented in the last row of Figure 3D, the classifier is less likely to decode water compared to the classical taste stimuli (Marascuilo’s test, Tables 1 and 2) and is more likely to decode sucrose and quinine compared to NaCl and citric acid (Marascuilo’s test, Tables 1 and 2). Altogether, these data indicate that ensembles of MD neurons recorded from active licking mice dynamically encode gustatory information up to 2 s after stimulus delivery. Considering that the neural activity in response to taste stimuli was recorded in the absence of overt differences in oromotor behaviors related to palatability (licking; Figs. 2C and 3B), the encoded information likely reflects the chemosensory discriminative aspects of the stimulus. In addition, taste coding showed a relatively ”late” onset - at least compared to the taste cortex (Bouaichi and Vincis, 2020) - with the time course of the averaged population decoding performance significantly rising above shuffled control within approximately 200 ms.

**Figure 3.**
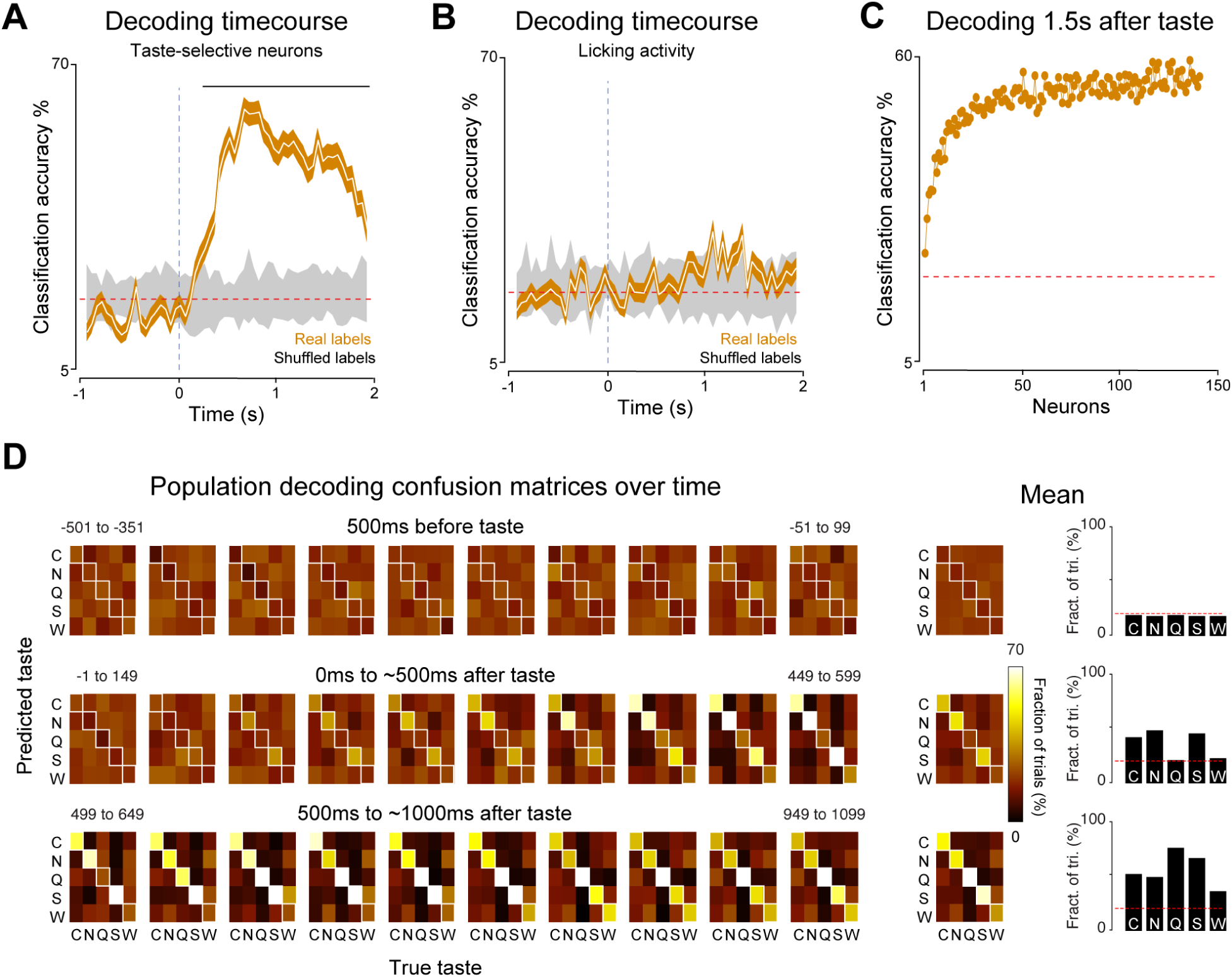
(A) Time course of decoding performance (white line) considering the population of taste-selective neurons. Gold-shaded area indicates the 1%ile to 99%ile range of the 20 times the decoder was run, each time using different training and testing splits (n = 20). Gray-shaded areas indicate the 1%ile to 99%ile range of the decoding performance over time after shuffling (20 times) stimulus labels for all trials. The horizontal black bar above the trace denotes bins when the classification accuracy significantly differed from the shuffled (permutation test, p *<* 0.05). (B) Time course of decoding performance (white line) considering the licking data extracted from the behavioral sessions from which the taste-selective neurons where recorded. Gold-shaded area indicates the 1%ile to 99%ile range of the 20 times the decoder was run, each time using different training and testing splits (n = 20). Gray-shaded areas indicate the 1%ile to 99%ile range of the decoding performance over time after shuffling (20 times) stimulus labels for all trials. (C) The mean accuracy of the decoder trained to discriminate the different gustatory stimuli (using a temporal window of 1.5 s after taste delivery) as the decoder gained access to progressively more neurons (gold dots). (D) Confusion matrices showing decoding performance for each gustatory stimulus in different 150-ms bins with 50-ms moving window temporal epochs around taste delivery (0 s). Color codes the classification accuracy, with bright hues indicating a higher fraction of correct trials. The main diagonal highlights the number of trials in which the classifier correctly assigned the taste stimulus (predicted taste) to its real category (true taste). On the rightmost panels the average decoding accuracy for three 500-ms temporal windows (-500 to 0 ms, top panels; 0 to 500 ms, middle panels; 500 to 1000 ms, bottom panels; 0 s taste delivery) shown both as confusion matrices and bar plots. In the bar plots, red dotted lines represent chance level.

**Table 1.** Marascuilo Multiple Comparison **Figure 3D middle panel** (“0ms to ∼500ms after taste”). The Marascuilo procedure was used to perform pairwise comparisons of multiple proportions after a chi-square test in Figure 3D. Here we report the pairwise differences between the proportions and their corresponding critical. The statistic was calculated for a significance level of *p <* 0.001. A difference is statistically significant if its value exceeds the critical range value. Sample proportions: *C* = 0.420, *N*= 0.482, *Q* = 0.211, *S* = 0.450, *W* = 0.229.

| Contrast | Difference | Critical_Range | Significance |
| --- | --- | --- | --- |
| C - N | 0.014 | 0.088 | Not Significant |
| C - Q | 0.248 | 0.081 | Significant |
| C - S | 0.147 | 0.085 | Significant |
| C - W | 0.161 | 0.086 | Significant |
| N - Q | 0.262 | 0.081 | Significant |
| N - S | 0.161 | 0.085 | Significant |
| N - W | 0.147 | 0.086 | Significant |
| Q - S | 0.101 | 0.079 | Significant |
| Q - W | 0.409 | 0.079 | Significant |
| S - W | 0.308 | 0.083 | Significant |

**Table 2.** Marascuilo Multiple Comparison **Figure 3D lower panel** (“500ms to ∼1000ms after taste”). The Marascuilo procedure was used to perform pairwise comparisons of multiple proportions after a chi-square test in Figure 3D. Here we report the pairwise differences between the proportions and their corresponding critical. The statistic was calculated for a significance level of *p <* 0.001. A difference is statistically significant if its value exceeds the critical range value. Sample proportions: *C* = 0.512, *N*= 0.498, *Q* = 0.760, *S* = 0.659, *W* = 0.351.

| Contrast | Difference | Critical_Range | Significance |
| --- | --- | --- | --- |
| C - N | 0.062 | 0.087 | Not Significant |
| C - Q | 0.209 | 0.079 | Significant |
| C - S | 0.030 | 0.087 | Not Significant |
| C - W | 0.191 | 0.080 | Significant |
| N - Q | 0.271 | 0.080 | Significant |
| N - S | 0.032 | 0.087 | Not Significant |
| N - W | 0.253 | 0.081 | Significant |
| Q - S | 0.239 | 0.080 | Significant |
| Q - W | 0.018 | 0.073 | Not Significant |
| S - W | 0.221 | 0.081 | Significant |

It is important to note that these results are obtained by administering taste stimuli at fixed and relatively low concentrations. This approach was primarily chosen to facilitate comparison with previous neural recordings in the mouse primary taste cortex (Levitan et al., 2019; Bouaichi and Vincis, 2020; Neese et al., 2022). Although the concentrations selected for each taste quality are suprathreshold, they are not equi-detectable. Therefore, differences in the MD neural activity may also result from changes in stimulus concentration. Concentration is a critical aspect of taste stimuli, and many studies have shown that perceptual/behavioral and neural responses to taste stimuli often correlate with concentration (Sadacca et al., 2012; Breza and Contreras, 2012; Spector et al., 2015; Wilson and Lemon, 2014; Fonseca et al., 2018). For these reasons, our next step was to explore if and how MD neurons integrate taste concentration information, focusing specifically on the taste qualities of sweet (sucrose) and salty (NaCl).

### Taste concentration processing in the mouse MD

To begin investigating how individual MD neurons represent sucrose concentration, we recorded individual units as described above. However, after habituation to head restraint, water-deprived mice were engaged in a task in which, this time, they had to lick six times on a spout to obtain a 3-µl drop of one of five peri- and suprathreshold sucrose concentrations (0.02 M, 0.04 M, 0.06 M, 0.1 M and 0.3 M)(Treesukosol et al., 2009; Spector, 2015), all presented at room temperature. Figure 4A shows the raster plots and PSTHs of two MD neurons that produced distinct responses to different sucrose concentrations. The responses of both neurons differed from the baseline spiking activity and showed a trend of increasing (Neuron #1) and decreasing (Neuron #2) firing rates with increasing stimulus concentration. Interestingly, monotonic positive and negative responses to changes in taste concentrations have been reported in other brain regions along the taste pathway (Sadacca et al., 2012; Yamamoto et al., 1984; Wilson and Lemon, 2014; Breza and Contreras, 2012; Fonseca et al., 2018). To quantify how many MD neurons respond selectively to sucrose concentrations, we used an approach similar to the one described above for taste quality. MD neurons were defined as concentration-selective if they exhibited a significant change in activity compared to baseline for at least one stimulus and responded differently to multiple stimuli. Our analysis revealed that, of the 154 MD neurons recorded, more than half (56%, 87/154) were sucrose concentration-selective. To further evaluate their responsiveness as a function of concentration, we normalized the evoked firing rate in a 1.5-s post-stimulus interval to the one evoked by the lowest concentration in our stimulus panel (0.02 M, Fig. 4B). As suggested by the example neurons in Figure 4A, the average spiking activity of these neurons was correlated with the increase in sucrose concentrations in a positive or negative monotonic manner (positive monotonic: R^2^ = 0.94, p-value = 0.0058; negative monotonic: R^2^ = 0.93; p-value = 0.00749; Fig. 4B). Importantly, this correlation was found to be independent of oromotor licking activity (Fig. 4B). When comparing the different types of neural responses, we did not observe significant differences in the proportion of neurons showing positive or negative monotonic responses (*χ*^2^ test for given probabilities: *χ*^2^ = 0.10345, df = 1, p-value = 0.7477; Fig. 4B). However, it is important to note that while the correlation between sucrose concentration and single neuron firing rate appears strong, this may be influenced primarily by the highest supra-threshold concentration 0.3 M. Indeed, when we reanalyzed the data excluding the highest concentration, the linear correlation was lost (positive monotonic: R^2^ = 0.64, p-value = 0.19; negative monotonic: R^2^ = 0.2, p-value = 0.49), suggesting that for the concentrations tested here, the main difference at the level of single-neuron response is driven by the highest sucrose concentration rather than a continuous gradient across all concentrations.

**Figure 4.**
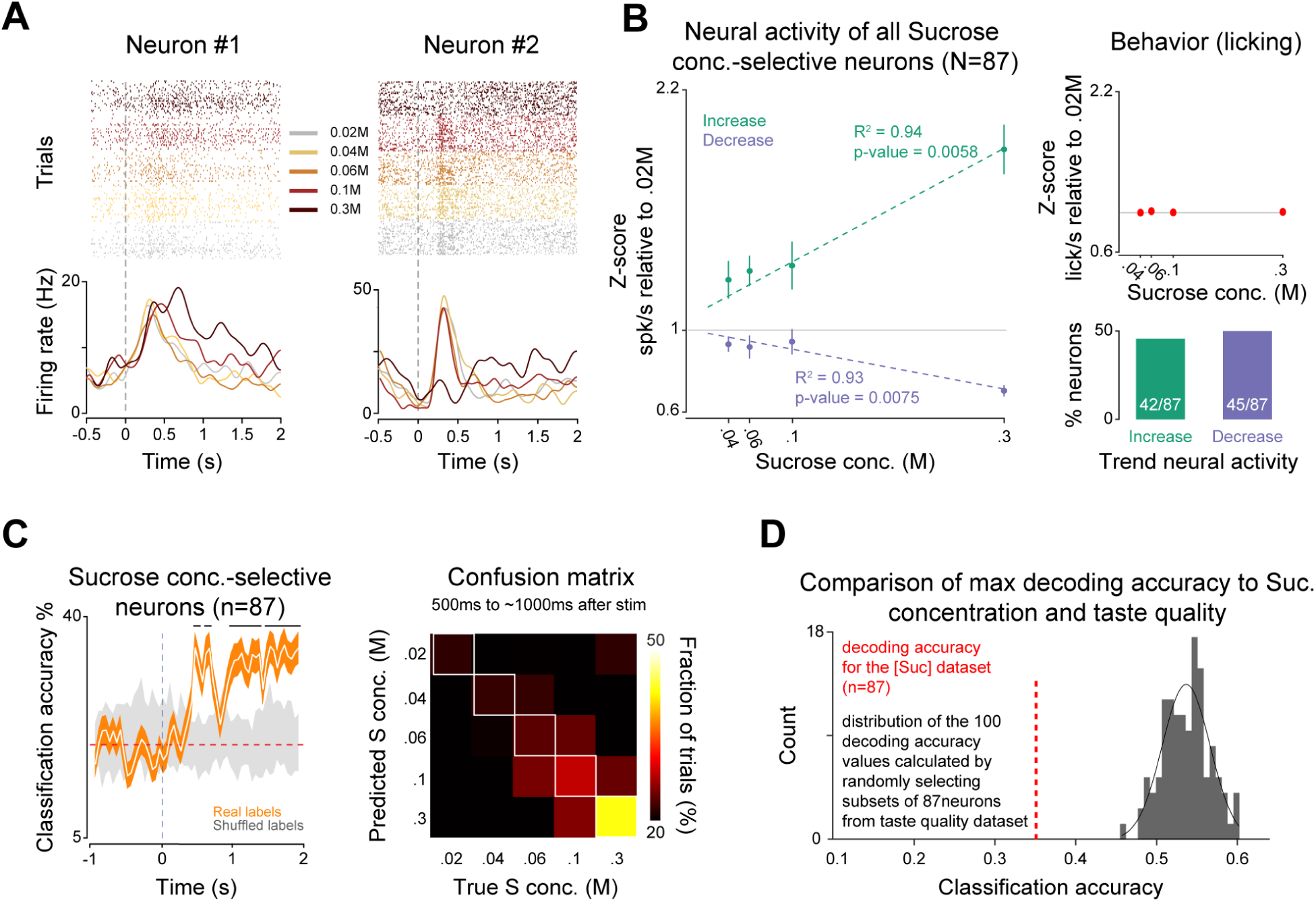
(A) Raster plots and PSTHs of two MD neurons showing sucrose concentration responses. Trials pertaining to different sucrose concentrations are grouped together (in the raster plots) and color-coded (both in the raster plots and PSTHs), with 0.02 M in gray, 0.04 M in yellow, 0.06 M in orange, 0.1 M in magenta, and 0.3 M in brown. (B) On the left panel, the average (circles) and standard error of the mean (vertical lines) of the firing rate of all neurons with sucrose concentration-related information that either increased (green) or decreased (purple) their activity as the sucrose concentration increased. Firing rate values are normalized to 0.02 M. The average Pearson’s R^2^ coefficients and associated p values are also shown for increases and decreases. On the top-right panel, the sucrose-independent lick rate (red circles) in the same time-window used for the quantification of spiking activity shown in the left panel. On the bottom-right panel, bar plot showing the fraction of neurons with sucrose concentration-related information that either increased (green) or decreased (purple) their activity as the sucrose concentration increased. (C) Left panel: time course of decoding performance (white line) considering the population of sucrose concentration-selective neurons. Gold-shaded area indicates the 1%ile to 99%ile range of the 20 times the decoder was run, each time using different training and testing splits (n = 20). Gray-shaded areas indicate the 1%ile to 99%ile range of the decoding performance over time after shuffling (20 times) stimulus labels for all trials. The horizontal black bars above the trace denote bins when the classification accuracy significantly differed from the shuffled (permutation test, p *<* 0.05). Right panel: confusion matrix showing decoding performance for each sucrose concentration in a 500-ms temporal window (500-1000 ms after stimulus delivery). Color codes the classification accuracy, with bright hues indicating a higher fraction of correct trials. The main diagonal highlights the number of trials in which the classifier correctly assigned the predicted sucrose concentration to its real category. (D) In black, the distribution of max decoding accuracy of taste quality obtained by randomly selecting 87 neurons from the taste quality dataset (n = 141, shown in Figures 2 and 3); the dotted vertical red line highlights the max decoding accuracy obtained from the sucrose concentration-selective neurons (n = 87).

We then wanted to investigate how well sucrose concentration was encoded at the level of population of MD neurons. To this end, we calculated the decoding performances using the neural population of sucrose concentration-selective neurons (Fig. 4C). The classification accuracy of the sucrose concentration-selective population exceeded the chance level and was significantly different from that of the shuffled control beginning more than 250 ms after stimulus delivery (permutation test, p-value *<* 0.05). The delayed onset of significant decoding appeared to be in accordance with the timing described for the onset of taste quality decoding (Fig. 3A). However, the decoding performance appeared to be qualitatively lower for sucrose concentration compared to taste quality. Although this comparison involves different populations of MD neurons, which could partly explain the observed difference, we can speculate as to why the average classification performance for sucrose concentration is lower than the one for taste quality. One reason could be that MD-evoked neural activity contains less information, particularly at lower sucrose concentrations, leading to lower average classification accuracy. This is supported by the confusion matrix shown in the right panel of Figure 4C; the classification performance for each concentration in a time window of 500 ms indicates that higher sucrose concentrations were better classified than lower ones. Another possibility is that the population of neurons that were selective for sucrose concentration was smaller than the group of neurons selective for taste, 87 and 141 neurons, respectively, which may explain the reduced classification. However, when we compared the maximum decoding accuracy for sucrose concentration with a distribution of maximum decoding accuracy for taste quality using subsets of 87 neurons, we found that the sucrose concentration performance lay below the 5^th^ percentile of the maximum decoding performance distribution for taste quality (Fig. 4D).

Next, we explored the extent to which MD neurons represent the concentration differences of another taste quality, salt. This time after mice had become accustomed to head restraint, water-restricted animals were trained on a licking task where they needed to lick six times on a spout to receive a 3-µl drop of one of five peri- and supra-threshold NaCl concentrations (0.02 M, 0.04 M, 0.07 M, 0.15 M, and 0.3 M)(Treesukosol and Spector, 2012; Spector, 2015). Figure 5A shows the raster plots and PSTHs of two MD neurons that produced NaCl concentration-selective responses. In this case, the responses of both neurons differed from the baseline spiking activity and showed a trend of increasing firing rates with increasing NaCl concentration.

**Figure 5.**
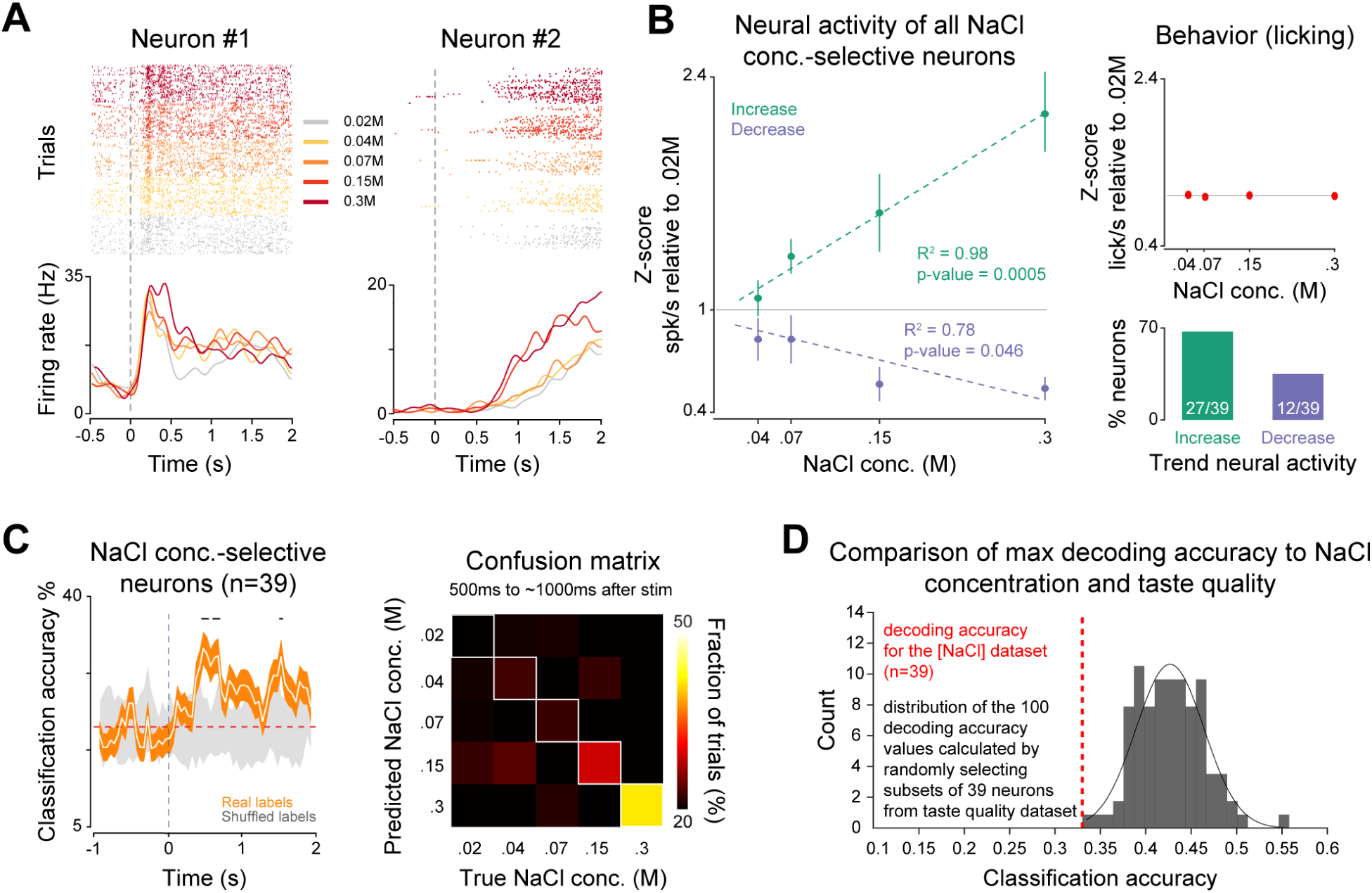
(A) Raster plots and PSTHs of two MD neurons showing NaCl concentration responses. Trials pertaining to different NaCl concentrations are grouped together (in the raster plots) and color-coded (both in the raster plots and PSTHs), with 0.02 M in gray, 0.04 M in yellow, 0.07 M in orange, 0.15 M in red, and 0.3 M in brown. (B) On the left panel, the average (circles) and standard error of the mean (vertical lines) of the firing rate of all neurons with NaCl concentration-related information that either increased (green) or decreased (purple) their activity as the NaCl concentration increased. Firing rate values are normalized to 0.02 M. The average Pearson’s R^2^ coefficients and associated p values are also shown for increases and decreases. On the top-right panel, the NaCl-independent lick rate (red circles) in the same time window used for the quantification of spiking activity shown in the left panel. On the bottom-right panel, bar plot showing the fraction of neurons with NaCl concentration-related information that either increased (green) or decreased (purple) their activity as the NaCl concentration increased. (C) Left panel: time course of decoding performance (white line) considering the population of NaCl concentration-selective neurons. Gold-shaded area indicates the 1%ile to 99%ile range of the 20 times the decoder was run, each time using different training and testing splits (n = 20). Gray-shaded areas indicate the 1%ile to 99%ile range of the decoding performance over time after shuffling (20 times) stimulus labels for all trials. The horizontal black bars above the trace denote bins when the classification accuracy significantly differed from the shuffled (permutation test, p *<* 0.05). Right panel: confusion matrix showing decoding performance for each NaCl concentration in a 500-ms temporal window (500-1000 ms after stimulus delivery). Color codes the classification accuracy, with bright hues indicating a higher fraction of correct trials. The main diagonal highlights the number of trials in which the classifier correctly assigned the predicted NaCl concentration to its real category. (D) In black, the distribution of max decoding accuracy of taste quality obtained by randomly selecting 39 neurons from the taste quality dataset (n = 141, shown in Figures 2 and 3); the dotted vertical red line highlights the max decoding accuracy obtained from the NaCl concentration-selective neurons (n = 39).

Our analysis revealed that, of the 84 MD neurons recorded, 46% were NaCl concentration-selective. Similarly to what was described for sucrose, we found that stimulus-evoked spiking activity was correlated with the increase in concentrations in a positive and negative monotonic manner (positive monotonic: R^2^ = 0.98, p-value = 0.0005; negative monotonic: R^2^ = 0.78, p-value = 0.0466; Fig. 5B) and was independent of oromotor licking. Similarly to what we described above for sucrose taste, we reanalyzed the data excluding the highest supra-threshold NaCl concentration. In this case, removing the highest concentration did not eliminate the correlation (positive monotonic: R^2^ = 0.96, p-value = 0.0196; negative monotonic: R^2^ = 0.93, p-value = 0.0333), indicating that for the NaCl concentration tested here, the firing properties of individual MD neurons remain correlated to the stimulus gradient even in the absence of the highest concentration. In addition, we found a higher proportion of neurons showing positive monotonic responses (*χ*^2^ test for given probabilities: *χ*^2^ = 5.7692, df = 1, p-value = 0.01631; Fig. 5B). The time course of the NaCl classification average across concentrations (Fig. 5C, *left panel*) showed the onset and accuracy similar to what was described for sucrose, with higher NaCl concentrations classified better than lower ones (Fig. 5C, *right panel*). Finally, when comparing decoding accuracy using subsets of neurons, NaCl concentration performance was also significantly lower, below the 1st percentile of the taste quality performance distribution (Fig. 5D).

Together, these data revealed that a substantial fraction of neurons in the mediodorsal thalamus represents sucrose and NaCl concentrations (and/or the “concentration” of water), with spiking activity that was correlated with the increase in stimulus concentrations in a positive and negative monotonic manner and independent of behavioral correlates of palatability (licking). However, our single-neuron analyses suggest that, for sucrose, this correlation is primarily driven by the highest concentration, whereas for NaCl, the relationship persists even after its removal. Furthermore, decoding analysis indicates that while individual MD neurons exhibit some degree of concentration selectivity, the discrimination of taste stimulus concentrations appears to be more robust at the population level, with higher concentrations being classified more accurately than lower ones. These findings support the idea that MD may contribute to the processing of sensory discriminative features of gustatory chemosensory stimuli. However, it is important to note that although the concentrations tested here span a relatively broad range of peri-threshold and supra-threshold values based on previous studies (Treesukosol and Spector, 2012; Spector, 2015), our mice were “passively” experiencing these stimuli without using this information to perform a behavioral task, such as detection or discrimination. Given this, the concentration-selective activity observed here, both at the single-neuron level and, more prominently, at the population level, likely represents only the tip of the iceberg in terms of the potential role of MD in concentration-selective gustatory processing. It is plausible that when concentration information becomes behaviorally relevant—especially in the peri-threshold range—MD neurons would exhibit even stronger and more selective responses, reflecting their role in task-driven sensory processing.

### MD processing of taste-specific expectation

Gustatory experiences are often perceived against the background of previous expectation. Cues of all sensory modalities frequently provide information predictive of taste outcomes and play a crucial role in shaping taste-driven behaviors (Gardner and Fontanini, 2014; Vincis and Fontanini, 2016; Samuelsen et al., 2012; Livneh et al., 2017; Kusumoto-Yoshida et al., 2015). Experiments in rats have highlighted MD as a brain region that responds to sensory cues associated with a specific behavioral outcome (Oyoshi et al., 1996; Kawagoe et al., 2007) and to the association of intraorally sourced odor and taste stimuli relevant to flavor (Fredericksen and Samuelsen, 2022). To determine how neurons in the mouse MD process information related to a taste-specific expectation, we relied on a two-cue taste association task. We trained mice (n = 5) to associate one auditory stimulus with the delivery of 0.3 M sucrose and another with the delivery of 0.03 M quinine (Fig. 6A). Two-second-long auditory cues (2 kHz and 12 kHz) were followed by a 0.5-s delay and by taste delivery (Fig. 6A). Cue-contingencies were counterbalanced across mice. To track associative learning, we analyzed the licking rate in anticipation of the two gustatory stimuli (Fig. 6B) by computing a *selectivity index* using a receiver operating characteristic (ROC) analysis (see Methods). The association was established for each individual animal when the *selectivity index* was negative (indicating higher lick rate for cue-sucrose association) and significant (p *<* 0.05) compared to the distribution of *pseudo selectivity indices* obtained after randomizing the trials. All mice reached the criterion within two weeks of training. After specific cue-taste associations were established, we performed neural recordings. Figure 6C illustrates four representative examples of MD neurons that were modulated by the presentation of auditory cues. Visual inspection of raster plots and PSTHs (Fig. 6C) indicated that, from a qualitative point of view, the activity of the MD neuron could be enhanced or suppressed by both auditory cues, irrespective of the valence of their associated outcome. It should be noted that cue responses consistently preceded the onset of licking. To quantify the fraction of thalamic neurons whose activity was modulated by different auditory stimuli, we analyzed changes in the firing rate that occurred before and after the onset of the cue (see Methods). Of all MD neurons recorded, an overwhelming majority (90.7%; 78/86; *χ*^2^ test for given probabilities: *χ*^2^ = 56.977, df = 1, p-value = 4.41e-14) were modulated by at least one auditory signal (Fig. 6D). Next, we analyzed whether and to what extent the activity of cue-responsive neurons provided information predictive of specific gustatory outcomes (as suggested by the example neurons shown in Fig. 6C) or represented general anticipatory signals. We addressed this issue using the following analysis. For each cue-responsive neuron, we compared the spiking activity evoked by the two different auditory stimuli. Briefly, the post-stimulus firing rate (0-2.5 s) was divided into five 500-ms bins, and a two-way ANOVA was used with “cue” and “time” as variables (Levitan et al., 2019; Bouaichi and Vincis, 2020; Jezzini et al., 2013). Cue-responsive neurons that had significantly different responses to the two tones in either the main effect “cue” or the “cue” × “time” interaction were defined as cue-selective (Fig. 6D). This method revealed that the majority of cue-responsive neurons (73%, 57/78) responded selectively to cues predicting different outcomes, while the remaining (27%) likely responded to general anticipatory signals (Fig. 6D; *χ*^2^ test for given probabilities: *χ*^2^ = 16.615, df = 1, p-value = 4.578e-05). In addition, the vast majority (80%, 46/57; Fig. 6E) of cue-selective neurons responded to cues predicting both outcomes with only a small fraction responding to only one auditory stimulus (20%, 11/57; *χ*^2^ test for given probabilities: *χ*^2^ = 21.491, df = 1, p-value = 3.555e-06). Among the latter, a similar number of neurons were modulated only by the sucrose-predicting or the quinine-predicting cue (54%, 6/11 for cue-quinine; 45%, 5/11 for cue-sucrose; *χ*^2^ test for given probabilities: *χ*^2^ = 0.090909, df = 1, p-value = 0.763). It is important to note that, according to the cue selectivity training paradigm, it was not possible to determine whether a given neuron responded to sucrose and/or quinine prior to training. While this does not invalidate the observed cue-selective responses, it is a factor to consider when interpreting cue selectivity. Future studies will further investigate how pre-existing taste responsiveness may influence cue-driven MD neural activity.

**Figure 6.**
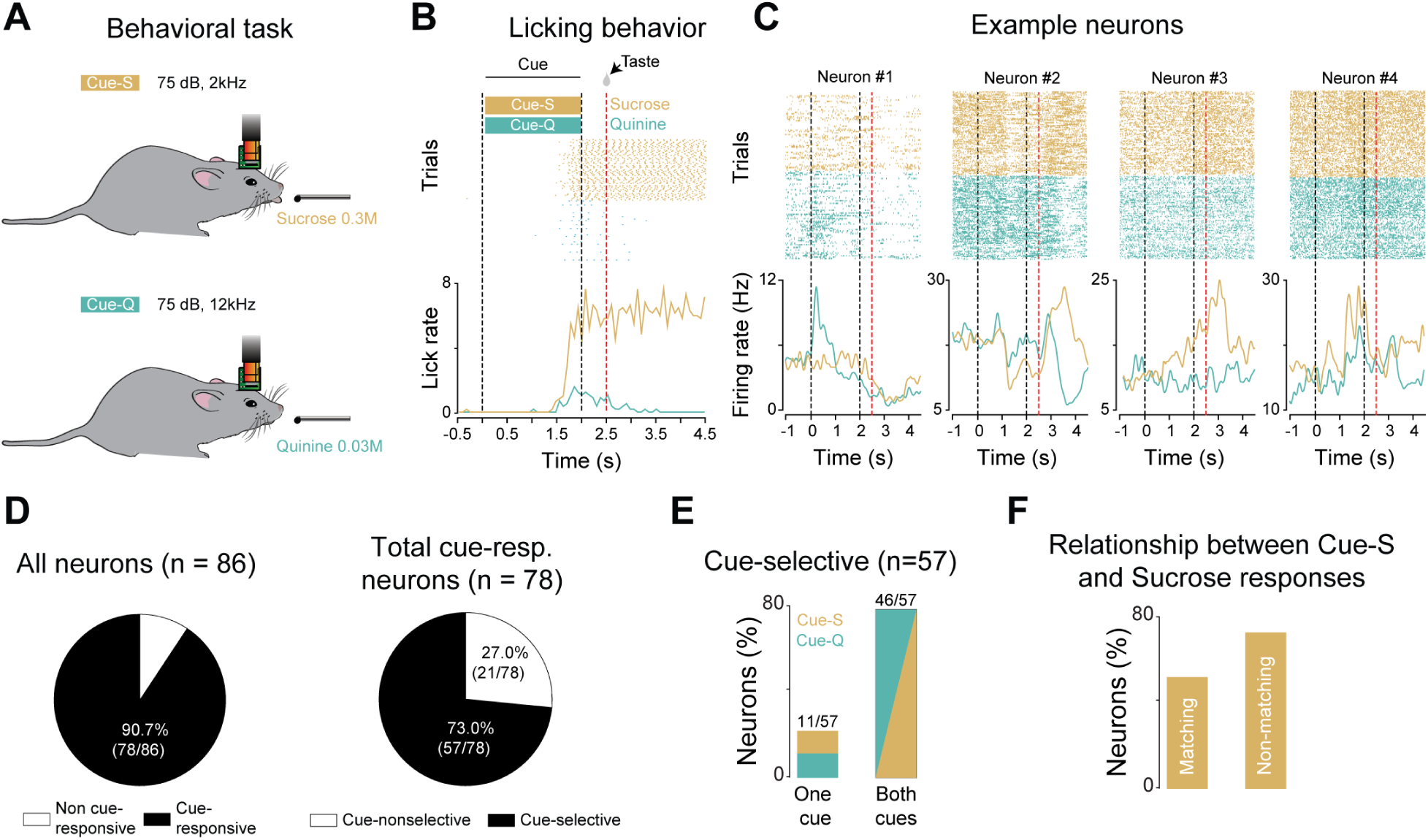
(A) Schematic representation of the behavioral task. Mice learned to associate specific auditory stimuli, tone 1 (gold) and tone 2 (green), with the delivery of two specific taste outcomes (sucrose and quinine, counterbalanced). (B) Raster plots and PSTHs of licking activity for a representative experimental session. In the raster plot, the colored areas indicate blocks of cue-sucrose (cue-S; gold) and cue-quinine (cue-Q; green) trials. The PSTHs show the average licking for cue-S and cue-Q trials in gold and green, respectively. The black dashed vertical lines mark the onset and offset of the auditory cue (0 s and 2 s, respectively), while the red dashed vertical line indicates taste delivery (2.5 s). (C) Raster plots and PSTHs for four MD cue-selective neurons. Raster plots and PSTHs are color coded, with activity evoked by cue-S and cue-Q shown in gold and green, respectively. Black dashed vertical lines indicate the onset and offset of the auditory cue (0 s and 2 s, respectively), while the red dashed vertical line indicates taste presentation (2.5 s). (D) Pie charts showing the proportion of MD cue-responsive (*leftmost chart*) and cue-selective (*rightmost chart*) neurons. (E) Fraction of cue-selective neurons responding to one or both auditory cues. (F) Fraction of cue-selective neurons modulated by cue-S and sucrose taste that exhibited matching responses (enhanced firing to both cue-S and sucrose taste; i.e., *Neuron#3 in panel C*) and non-matching responses (enhanced firing to cue-S but suppressed to sucrose taste, or vice versa; i.e., *Neuron#2 in panel C*).

Next, we performed a series of analyses to first determine whether cue-selective neurons also respond to the taste outcome and, second, to evaluate if the responsiveness of a neuron to a specific cue is related to its response to the taste. An important caveat must be recognized here. The successful learning of the cue-taste association task led to reduced anticipatory licking for cue-Q, resulting in very few trials with quinine consumption. As a consequence, modulation in the firing rate after “quinine” could result from movements in the mouth and snout associated with avoiding licking to avoid consumption. For this reason, we restricted these analyses to sucrose trials. Among sucrose cue-selective neurons, 56% (29/51) were also selectively modulated by the sucrose taste. We then performed an additional analysis to examine the relationship between their response to cue-S and their response to sucrose taste, specifically assessing whether the “sign” of neural modulation (enhancement or suppression) was consistent between both conditions (Fig. 6F). The results revealed that among MD neurons selectively modulated by cue-S, similar proportions exhibited matching responses (enhanced firing to both cue-S and sucrose taste) and non-matching responses (enhanced firing to cue-S but suppressed to sucrose taste, or vice versa) (*χ*^2^ test for given probabilities: *χ*^2^ = 0.31034, df = 1, p-value = 0.5775) (Fig. 6F).

Together, these results show that individual neurons in the mouse MD can respond selectively to cues anticipating taste outcomes. In addition, cue-selective neurons appear to preferentially represent information from cues predicting both rewarding and aversive taste, and some cue-selective neurons have different response patterns to the cues and the predicted outcomes.

### Population decoding of cue-taste expectation by the MD

After characterizing the profile of cue-related responses in single neurons, we focused our attention on neural activity at population level. Thus, to further evaluate how the MD network encodes specific expectations relevant to guiding consummatory behavior, we computed the decoding performances in the population of cue-responsive neurons. The time course of the classification average is shown in Figure 7A. The decoding performance of the cue-selective population (Fig. 7A) showed an early onset, with a classification accuracy above the chance level and significantly different from that of the control (i.e., shuffled data) from the first bin after cue onset (0–150 ms; permutation test, p *<* 0.05). As an additional control, the same analysis was performed using licking instead of spiking data (blue dotted line, Figure 7A). In particular, the decoding with neural data showed an onset that occurred seconds before the decoding with licking data, indicating that neural responses, especially those in the first 1.5 s after the onset of the auditory stimulus, precede the licking activity. Furthermore, the classification accuracy of the cue-selective population significantly differed from that of the shuffled control for the entire 5-s window including cue and taste-outcome (Fig. 7A). This suggests that cue-selective MD neurons are capable of encoding both anticipatory and taste-outcome information.

**Figure 7.**
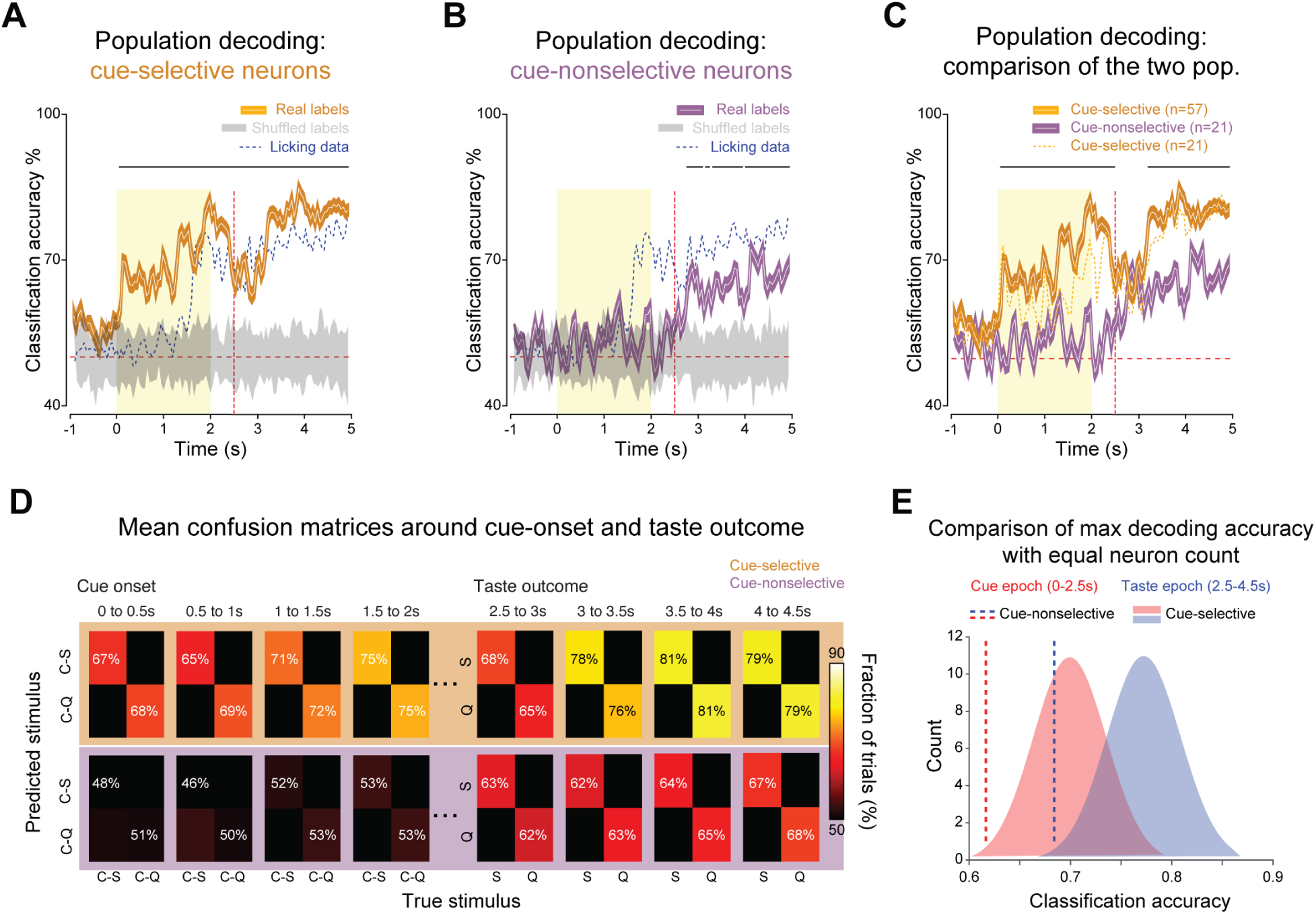
(A) Time course of decoding performance (white line) considering the population of cue-selective neurons. Gold-shaded area indicates the 1%ile to 99%ile range of the 20 times the decoder was run, each time using different training and testing splits (n = 20). Gray-shaded areas indicate the 1%ile to 99%ile range of the decoding performance over time after shuffling (20 times) stimulus labels for all trials. The horizontal black bar above the trace denotes bins when the classification accuracy significantly differed from the shuffled (permutation test, p *<* 0.05). Dotted blue line represents the time course of decoding performance when licking data were used. Yellow box indicates time of auditory cue presentation. Dotted horizontal red line indicates chance level (50%); dotted vertical red line indicates time of taste outcome delivery. (B) Time course of decoding performance (white line) considering the population of cue-nonselective neurons. Purple-shaded area indicates the 1%ile to 99%ile range of the 20 times the decoder was run, each time using different training and testing splits (n = 20). Gray-shaded areas indicate the 1%ile to 99%ile range of the decoding performance over time after shuffling (20 times) stimulus labels for all trials. The horizontal black bars above the trace denote bins when the classification accuracy significantly differed from the shuffled (permutation test, p *<* 0.05). Dotted blue line represents the time course of decoding performance when licking data were used. Yellow box indicates time of auditory cue presentation. Dotted horizontal red line indicates chance level (50%); dotted vertical red line indicates time of taste outcome delivery. (C) Time course of population decoding performance over time by the cue-selective (gold) and cue-nonselective neurons. The horizontal black bars above the trace denote bins when the classification accuracy significantly differed between the two populations (permutation test, p *<* 0.05). Dotted gold line represents the time course of decoding performance with a population of cue-selective neurons composed of 21 randomly selected cells. Yellow box indicates time of auditory cue presentation. Dotted horizontal red line indicates chance level (50%); dotted vertical red line indicates time of taste outcome delivery. (D) Confusion matrices showing decoding performance during different 500-ms temporal epochs around cue onset (0 s) and taste-outcome delivery (2.5 s) for the cue-selective (gold background) and cue-nonselective (purple background) populations. Color codes the classification accuracy, with bright hues indicating a higher fraction of correct trials. The main diagonal highlights the number of trials in which the classifier correctly assigned the stimulus (predicted stimulus) to its real category (true stimulus). (E) In red and blue, the distribution of max decoding accuracy during cue presentation (Cue epoch, 0-2.5 s; red) and after taste outcome (Taste epoch, 2.5-4.5 s; blue) obtained by randomly selecting 21 neurons from the cue-selective neuron population; the dotted vertical red and blue lines highlight the max decoding accuracy during cue presentation (Cue epoch, 0-2.5 s; red) and after taste outcome (Taste epoch, 2.5-4.5 s; blue) obtained from cue-nonselective neuron population (n = 21).

Next, we switched our focus to the population of cue-nonselective neurons, those MD neurons whose firing activity evoked by auditory cues differed from baseline but was similar between the two taste-predicting stimuli. As expected, the decoding performance of the cue-nonselective neurons showed that its population activity did not represent information about the auditory cues. However, the classification accuracy of the cue-nonselective neuron population exceeded the chance level and differed significantly from that of the control (i.e., shuffled data) beginning hundreds of milliseconds after taste outcome delivery (Fig. 7B; permutation test, p *<* 0.05). Thus, the cue-nonselective population of MD neurons may primarily process information related to actual taste outcomes rather than anticipatory auditory cues.

We then compared the classification accuracy of the two populations (Fig. 7C). As expected, the decoding performance of the cue-selective population differed significantly from that of the cue-nonselective population from the first bin after cue onset and continuing until taste outcome delivery (permutation test, p *<* 0.05). This can also be observed from a qualitative analysis of the confusion matrices shown in Figure 7D, where the classification performances for each stimulus are represented. The significant difference in classification accuracy between the two populations was briefly disrupted after taste outcome delivery (Fig. 7C-D), where both populations of MD neurons encoded taste outcome information with similar accuracy. The brief disruption in the significant difference after taste delivery could reflect a transient period in which both populations of neurons are equally capable of encoding the presence or absence of fluid. However, the significant difference in decoding performance reemerged for the remaining temporal window, highlighting that cue-selective neurons better encode sucrose taste information compared to cue-nonselective neurons (Fig. 7C-D). This is consistent with the time course of taste quality decoding shown in Figure 3 where the taste classification average showed a delayed onset _200 ms after taste delivery. It is important to note that from a quantitative point of view, the differences in the time course of the two populations do not appear to depend on the number of neurons forming each population (Fig. 7C, see dotted gold line). However, to further evaluate whether the decoding differences between the two populations during the cue period and, more importantly, during taste outcomes were the result of differences in population sizes, we performed an additional analysis. We compared the maximum decoding accuracy for the cues and taste outcomes of the cue-nonselective neurons with a distribution of the maximum decoding accuracy using subsets of 21 cue-selective neurons (Fig. 7E). The decoding accuracies of the cue-nonselective populations lay below the 5^th^ percentile of the decoding performance distribution for cue-selective neurons (Fig. 7E), suggesting that differences in encoding capabilities cannot be exclusively explained by the different population sizes.

Together, these results suggest that cue-selective neurons in the MD network are specifically tuned to auditory cues that predict different outcomes, allowing early and precise encoding of these signals. Further, cue-selective neurons also reliably encoded taste outcomes, demonstrating their dual role in both anticipatory and outcome processing. In contrast, cue-nonselective neurons responded more generally and only showed decoding performance - though not as accurately as cue-selective neurons - after the actual taste outcome was delivered, indicating their involvement in processing the outcome rather than predicting it. In summary, these results highlight the role of MD neurons in anticipatory encoding, which is crucial to guide behavior based on expected outcomes.

## Discussion

Although previous studies have demonstrated the role of MD in experience-dependent olfactory processing (Courtiol and Wilson, 2014; Plailly et al., 2008; Eichenbaum et al., 1980), recent findings in rodents suggest that MD is also involved in oral chemosensory processing. Specifically, after odor-taste associations, MD neurons have been shown to represent taste information, integrating these signals in a way that influences consummatory behavior and sensory attention (Fredericksen and Samuelsen, 2022; Gartner and Samuelsen, 2024). Considering that taste-odor associations can shape neural activity in chemosensory-processing regions (Vincis and Fontanini, 2016; Maier et al., 2023), the first question we explored in this study is whether MD processes gustatory signals solely within the framework of learned associations or if it can encode the sensory features of a taste stimulus independently of overt prior experience. Our findings provide novel evidence supporting the latter, demonstrating that MD neurons encode both taste identity and concentration when tastants are sampled by active licking, even in the absence of explicit associations with other oral stimuli. This expands our understanding of MD’s role in chemosensory processing, revealing its capacity to represent gustatory information beyond learned taste-odor associations. Using silicon probe recordings, we found that neurons in the medial subregion of mouse MD, which is reciprocally connected to GC (Fig. 1), respond in a broadly tuned fashion to different taste qualities and, at the population level, can accurately and dynamically encode both the identity and concentration of gustatory signals. Importantly, our results indicate that the taste-evoked responses in MD appear to be independent of variations in licking behavior. Considering that in rodents, palatability is operationally defined by changes in oromotor responses (Spector and St. John, 1998), this suggests that MD activity may primarily reflect the qualitative discrimination of taste stimuli rather than their hedonic value. This raises the question of how taste-responsive neurons in MD compare with those in GC, the amygdala, and the prefrontal cortex—regions reciprocally connected to MD (Ahmed and Paŕe, 2023; Mitchell and Chakraborty, 2013) and previously studied for their role in taste processing (Katz et al., 2001; Sadacca et al., 2012; Grossman et al., 2008; Bouaichi and Vincis, 2020; Jezzini et al., 2013). Although the overall accuracy of taste quality coding and breadth of tuning are largely similar, the temporal profile of taste responses differs. Specifically, a qualitative evaluation of the decoding performance suggests that MD encodes taste information more slowly than GC and the basolateral amygdala (BLA) (Grossman et al., 2008; Bouaichi and Vincis, 2020), but faster than the prefrontal cortex (Jezzini et al., 2013), supporting the idea that MD might act as an intermediate processing hub between sensory and higher-order areas within the gustatory system. From a broader perspective, this leads to the question of the functional role of MD within the taste network for processing neural information relevant for consummatory behavior. An admittedly over-simplistic possibility is that MD mainly inherits gustatory information from these inputs, consistent with the broader principle that similar neural representations across interconnected brain regions can provide functional redundancy to enhance the robustness of neural processing (Johnston and Freedman, 2023). An alternative and perhaps more accurate possibility is that MD does not exclusively “duplicate” taste processing occurring in GC and the amygdala but instead provides an additional computational layer where gustatory chemosensory signals are integrated with orally sourced olfactory cues and potentially modulated by associative and experience-dependent factors (Fredericksen and Samuelsen, 2022). Recent work has shown that pharmacological inactivation of the rat’s MD alters consummatory behaviors related to the hedonic value of previously associated odor-taste mixture stimuli and affects sensory attention (Gartner and Samuelsen, 2024). In this framework, MD taste responses in the absence of prior experience - as the ones observed in our study - may serve as a functional footprint where gustatory features from GC and the amygdala could be integrated with orally sourced odor information. Through associative learning, memory, and decision-making circuits, the MD may thus contribute to establishing flexible representations of chemosensory stimuli relevant to consummatory behaviors (Gartner and Samuelsen, 2024). Future research investigating MD activity in learning and task-dependent conditions, particularly in relation to its reciprocal connections with GC and the amygdala, will be crucial for understanding how this thalamic hub shapes adaptive chemosensory processing and orchestrates neural dynamics relevant to consummatory behaviors.

Multiple studies have indicated that exteroceptive signals associated with food-related stimuli modulate consummatory behavior, drive neural activity, and alter taste coding in the gustatory-related brain regions (Spence, 2015; Gardner and Fontanini, 2014; Livneh et al., 2017; Samuelsen et al., 2012; Liu and Fontanini, 2015). Here, we investigated how MD, a region often studied for its associative characteristics (Mitchell and Chakraborty, 2013), responds to auditory signals that selectively predict a rewarding (sucrose) or an aversive (quinine) gustatory outcome, and - at least in the case of sucrose-predicting cues - how these responses relate to the neural activity evoked by the predicted taste. Our results show that, after their association with taste, auditory cues gain selective control of the activity of a large majority of neurons recorded in the MD (Fig. 6). Moreover, in our study, the vast majority of MD cue-selective neurons (e.g., those whose cue-evoked firing rates significantly differed between cue-S and cue-Q) responded to both types of predictive signals without a bias toward reward-associated cues, unlike previous reports where MD neurons predominantly encoded rewarding outcomes (Kawagoe et al., 2007; Oyoshi et al., 1996). These responses could reflect either the expectation of specific taste qualities (e.g., sucrose vs. quinine) or a broader anticipation of reward presence vs. absence. While our task does not fully disambiguate these models, the presence of taste-selective responses support the idea that MD could encode taste identity predictions rather than simply signaling reward availability. Regardless of this distinction, it is interesting to speculate on how MD fits within the broader neural circuit governing taste expectation. Given its reciprocal connections with both GC and BLA—regions central to current models of expectation-related taste coding (Kusumoto-Yoshida et al., 2015; Gardner and Fontanini, 2014)—MD is well-positioned to influence how predictive cues shape taste perception and consummatory behaviors. The decoding performance of the cue-selective population (Fig. 7A) showed a rapid onset, comparable to that observed in GC (Gardner and Fontanini, 2014). This similarity raises a key question: does MD, through its interactions with the frontal cortices and amygdala, drive the emergence of cue-related activity in GC (Schoenbaum and Roesch, 2005; Samuelsen et al., 2012)? Alternatively, are parallel mechanisms at play, with MD encoding predictive cue signals independently of these regions? Understanding whether MD actively drives cue-related activity in GC and BLA or whether it serves as a complementary circuit processing expectation signals in parallel will be crucial for refining our models of taste-expectation processing. Another interesting point to consider is whether MD neurons inherently differentiate between two auditory stimuli or if auditory signals gain selective control over MD activity as a function of learning. Given prior research, it is reasonable to speculate that the latter is more likely. On one hand, while there is no strong evidence that MD receives direct auditory inputs, indirect pathways from the prefrontal cortex (Parnaudeau et al., 2013), BLA (Groenewegen et al., 1999), and mesencephalic nuclei (MN) (Shin et al., 2023) could convey auditory information. However, unlike bottom-up auditory pathways, these indirect routes may provide MD with associative, experience-dependent signals or sound-induced arousal rather than purely sensory input. On the other hand, MD has been shown to be heavily involved in associative learning (Mitchell and Chakraborty, 2013), and the appearance of cue responses have been shown to be strongly dependent on learning (Gardner and Fontanini, 2014; Samuelsen et al., 2012; Vincis and Fontanini, 2016).

Previous studies suggested that, at least in sensory brain regions, expectations can trigger the anticipatory activation of stimulus-specific representations (Zelano et al., 2011). For instance, in the GC, neurons that respond to predictive cues often exhibit a corresponding pattern of activity upon taste delivery, where neurons showing cue-evoked enhancement or suppression in firing rate tend to respond to sucrose with the same sign, suggesting a strong link between expectation and sensory processing (Gardner and Fontanini, 2014). Our data indicate that, unlike in GC, MD neurons do not exhibit a systematic cue-taste relationship, as cue-selective neurons were equally likely to show matching responses (enhanced firing to both cue-S and sucrose taste) or non-matching responses (enhanced firing to cue-S but suppressed to sucrose taste, or vice versa). Interestingly, a lack of systematic alignment of cue-taste has been observed in the nucleus accumbens (NAc) (Roitman et al., 2005) - another region of the brain that is reciprocally connected to the medial portion of the MD (Erro et al., 2000; Groenewegen et al., 1999) - suggesting that these regions are also capable of encoding predictive signals differently than sensory outcomes. It is tempting to speculate that given the participation of MD in the flexible updating of sensory-motor representations (Parnaudeau et al., 2013; Mitchell and Chakraborty, 2013; Gartner and Samuelsen, 2024), the observed variability in cue-taste relationships observed here may reflect a broader mechanism in which MD also contributes to context-dependent modulation of taste-guided behaviors, rather than rigidly encoding stimulus-specific predictions. In summary, while the mediodorsal thalamus is often overlooked in the context of the taste neural circuit, our findings imply that this assumption may need to be revisited. Our results show that MD neurons dynamically encode gustatory chemosensory signals and selectively respond to cues that predict rewarding and aversive taste outcomes, which are important in driving consummatory decisions. Future studies involving neural manipulation in behaving animals will reveal the full spectrum of the role of the MD in gustatory perception.

## Acknowledgements

This research was supported by grant number R01 DC-019326 from the National Institute on Deafness and Other Communication Disorders to RV.

